# *Bacillus velezensis* EU07 suppresses *Fusarium graminearum* via transcriptomic reprogramming

**DOI:** 10.1101/2025.10.10.681611

**Authors:** Ömür Baysal, Catherine Jimenez-Quiros, Birsen Cevher-Keskin, Mahmut Tör

## Abstract

*Fusarium graminearum*, the causal agent of Fusarium head blight, is a devastating pathogen of cereals worldwide. Biological control using *Bacillus* species has emerged as a sustainable strategy to suppress this pathogen, but the molecular basis of antagonism remains poorly understood. Here, we investigated the interaction between *Bacillus velezensis* EU07 and *F. graminearum* strain K1-4 through morphological assays and RNA-seq profiling. Microscopy revealed severe hyphal distortions including swelling and branching abnormalities, following exposure to EU07 cell pellets. Transcriptomic analysis after 6 h of treatment identified 1,264 differentially expressed genes (DEGs), with 732 downregulated and 532 upregulated. Genes encoding secondary metabolite biosynthesis enzymes, including trichothecene (TRI) cluster genes, cytochrome P450s, and transporters, were strongly repressed. Key metabolic pathways, such as amino acid catabolism and mitochondrial transporters (e.g., 2-oxoglutarate/malate carrier protein), also showed reduced expression. Conversely, genes associated with oxidative stress responses, detoxification, and membrane transport were induced, reflecting a compensatory survival strategy. These results demonstrate that EU07 disrupts *F. graminearum* both morphologically and at the transcriptional level, suppressing virulence-associated pathways while triggering stress adaptation. This dual impact highlights *B. velezensis* EU07 as a promising biocontrol agent and provides candidate fungal genes for targeted RNAi-based crop protection strategies.

## Introduction

Fusarium head blight (FHB), primarily caused by *Fusarium graminearum*, is a major disease of cereal crops worldwide. It leads not only to significant yield losses but also to contamination of grain with mycotoxins, such as deoxynivalenol (DON), which pose serious threats to food safety, and human and animal health (Wegulo 2012). Traditional approaches to manage FHB have relied heavily on chemical fungicides and resistant cultivars. However, the emergence of fungicide resistance, increasing regulatory restrictions on chemical inputs, and the limited protection conferred by resistant varieties emphasize the urgent need for sustainable alternative disease management strategies (Lee et al., 2023).

Biological control agents (BCAs), particularly beneficial microorganisms, offer an environmentally friendly and sustainable strategy for managing FHB (Gao et al., 2016; Zubair et al., 2021). These agents suppress pathogens through various mechanisms, including antibiosis, competition for space and nutrients, and the induction of systemic resistance in host plants (Blake et al. 2021). Compared to chemical fungicides, BCAs are generally biodegradable and less likely to induce resistance in pathogens.

Within this group, *Bacillus* species have shown particular promise due to their ability to form endospores, produce a wide array of secondary metabolites, and survive under a range of environmental conditions (Abdel-Aziz et al., 2017).

Species such as *B. subtilis*, *B. amyloliquefaciens*, *B. licheniformis*, and *B. pumilus* have demonstrated efficacy against various phytopathogens (Karačić et al., 2024). Their antagonistic activity is largely attributed to the production of antifungal lipopeptides such as iturins, fengycins, and surfactins that inhibit fungal spore germination, disrupt hyphal integrity, and interfere with fungal signalling pathways (Ongena et al., 2008). These compounds not only inhibit fungal growth but also trigger systemic resistance mechanisms in host plants (Ongena et al., 2007). Some strains also produce volatile organic compounds and enzymes capable of degrading fungal cell walls or detoxifying fungal metabolites like fusaric acid (Smaoui et al., 2023; Wadhwa et al., 2024).

Interestingly, recent studies have pointed to the enhanced efficacy of microbial consortia over single-strain applications (Comite et al., 2021; Nunes et al., 2024; Pérez-Moncada et al., 2024). Carefully selected bacterial-fungal consortia can broaden the spectrum of disease suppression and increase the reliability of biocontrol under variable environmental conditions. Arbuscular mycorrhizal fungi (AMF), for example, have been shown to synergize with bacteria in activating plant defense pathways and modulating rhizosphere interactions (Whipps 2001; Kashyap et al., 2024; Weisany, 2024). Furthermore, root-colonizing strains of *Bacillus*, particularly those isolated from plant-associated niches, tend to exhibit superior biocontrol potential and rhizosphere competence. These strains are increasingly being integrated into next generation bioformulations.

Despite these advances, interactions between BCAs and target pathogens remain complex and occasionally contradictory. While *Bacillus* strains generally suppress *F. graminearum* growth and mycotoxin production, certain metabolites such as bacillomycin D have been reported to inadvertently stimulate DON production in some cases (Gu et al., 2017). These observations highlight the need for mechanistic investigations that move beyond phenotypic assays to explore the molecular and transcriptional dynamics of pathogen-antagonist interactions.

Transcriptomic approaches have proven instrumental in unraveling these dynamics. RNA-seq analyses have revealed that pre-treatment of plants with *Bacillus* strains leads to the upregulation of jasmonic and salicylic acid pathway genes in the host, as well as pathogenesis-related proteins such as PR-1 and PR-10, upon pathogen challenge (Le Henanff et al., 2009; Rabari et al., 2023; Gebarowska et al., 2023; Gupta et al., 2024; Zhang et al., 2025). On the microbial side, transcriptomic profiling has revealed major shifts in *Bacillus* metabolic and regulatory networks during antagonistic interactions, particularly in secondary metabolite biosynthesis, redox balance, and nutrient metabolism (Medeiros et al., 2011; Wahab et al., 2023). For instance, *B. velezensis* LZN01 showed significant transcriptional changes when optimized for antifungal activity against *Fusarium oxysporum*, with hundreds of genes involved in primary and secondary metabolism being differentially expressed (Hu et al., 2024; Assena et al., 2024).

In our previous work, we identified and characterized the *B. velezensis* strain EU07, which exhibited strong antagonistic activity against *F. graminearum* K1-4 both *in vitro* antagonism and *in planta* infection assays (Jimenez-Quiros et al., 2022). Beyond its antifungal efficacy, EU07 also promoted plant growth, suggesting its potential as a dual function biopesticide and plant growth-promoting rhizobacterium (PGPR). Comparative genomic and proteomic analyses placed EU07 within the *B. subtilis* species complex and highlighted its unique protein expression profile relative to commercial strains, indicating distinctive metabolic and biocontrol capabilities (Baysal et al., 2013; Nikolaidis et al., 2022).

In the current study, we extend this work by using RNA-seq to profile transcriptomic changes in *F. graminearum* K1-4 in response to EU07 treatment. Our aim was to identify differentially expressed genes (DEGs) associated with key metabolic, regulatory, and virulence pathways. By characterizing these transcriptional responses, we seek to gain mechanistic insights into the antifungal activity of EU07 and to pinpoint fungal genes that may serve as candidates for RNA interference (RNAi)-based control strategies. Understanding how EU07 suppresses *F. graminearum* at the molecular level will inform the design of more targeted and effective biocontrol products.

## Results

### Optimising conditions for transcriptomic profiling of *F. graminearum* treated with *B. velezensis* EU07

We conducted a preliminary experiment to identify treatment conditions under which *F. graminearum* K1-4 (*Fg*-K1-4) shows a reproducible physiological response to *B. velezensis* EU07, suitable for transcriptomic profiling.

An exploratory assay was designed to test the effect of different EU07-derived treatments on fungal morphology. Five conditions were compared: (i) untreated control, (ii) LB broth only, (iii) whole EU07 culture broth, (iv) cell-free culture supernatant (centrifuged and filtered), and (v) EU07 bacterial pellet washed and resuspended in sterile water. All treatments were applied to fungal cultures under identical conditions, each with three biological replicates.

Morphological changes were assessed 48 h post-treatment. In untreated controls (Figure 1A), fungal hyphae appeared normal. In contrast, LB broth and all EU07-derived treatments (Figure 1B–E) induced pronounced alterations, including localized swelling and rounded structures along the hyphae, indicative of stress or disrupted growth.

**Figure 1.**
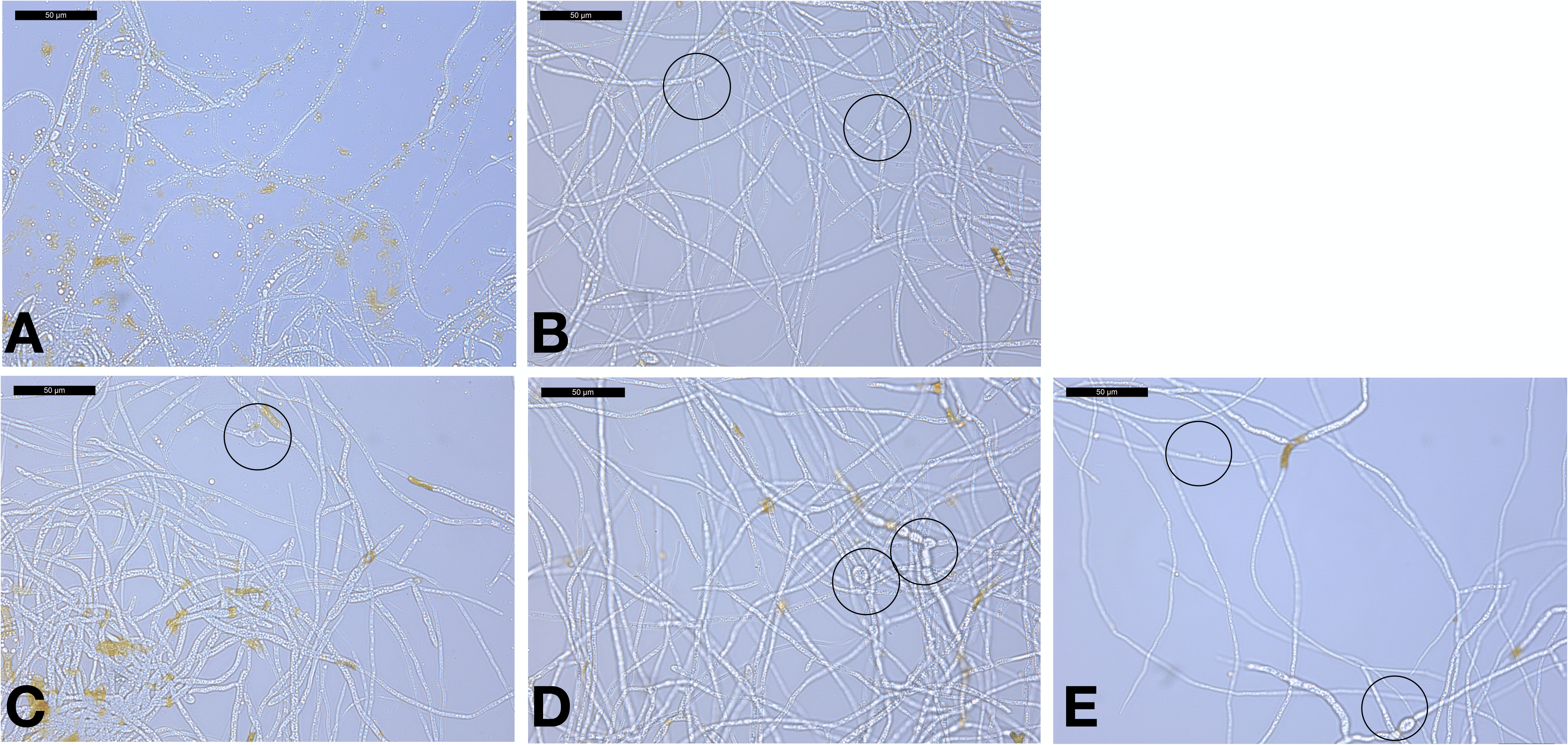
Morphological effects of *B. velezensis* EU07-derived treatments on *F. graminearum*. Cultures were treated under four different conditions: **A)** Control-no treatment, **B)** LB broth only, **C)** Whole EU07 culture broth, **D)** Cell-free EU07 supernatant (centrifuged and 0.22 µm-filtered), and **E)** EU07 bacterial pellet washed and resuspended in sterile water. Scale bars = 50 µm. Circles indicate thick, rounded hyphal regions.

Among the tested conditions, washed EU07 cells resuspended in sterile water (E) were selected for subsequent transcriptomic analysis. This condition (i) consistently elicited a clear morphological response and (ii) enabled direct bacterial–fungal interactions without confounding effects from the components of LB broth.

This pilot study established reproducible and biologically relevant conditions for downstream transcriptomic profiling, providing a robust framework for examining fungal gene expression during interaction with bacterial cells.

### DEGs in F. graminearum in response to B. velezensis EU07

RNA sequencing of the six samples generated between 12.4 and 19.0 million clean reads per library (Supplementary Table 1). The proportion of effective reads exceeded 97% across all samples, with an error rate consistently below 0.05%. Quality assessment showed that more than 94% of bases had a Phred quality score above Q30, while the average GC content was approximately 52%. These metrics confirm that the datasets are of high quality and suitable for downstream transcriptomic analyses.

Transcriptomic analysis revealed significant differences in gene expression in *F. graminearum* exposed to EU07 metabolites (Supplemental Table 2). A number of genes were significantly downregulated compared to control conditions. Gene Set Enrichment Analysis (GSEA) indicated consistent suppression of genes across several functional categories (Table 1).

**Table 1.** Enriched pathways among DEGs in *F. graminearum* exposed to *B. velezensis* EU07 metabolites.

| Direction | adj. P-value | nGenes | Enriched pathway/domain |
| --- | --- | --- | --- |
| Downregulated | $1.2 \times 10^{-3}$ | 5 | RmlC-like cupin domain superfamily |
| Upregulated | $4.8 \times 10^{-4}$ | 3 | Amino acid transporter, transmembrane domain |
| Upregulated | $1.0 \times 10^{-2}$ | 2 | Amino acid permease / SLC12A domain |

Heatmap analysis highlighted a clear transcriptomic shift: genes normally downregulated under control conditions were upregulated following exposure to EU07 metabolites, while those typically upregulated were suppressed (Figure 2). This pattern suggests a complex regulatory response, possibly involving compensatory or stress-adaptive mechanisms.

**Figure 2.**
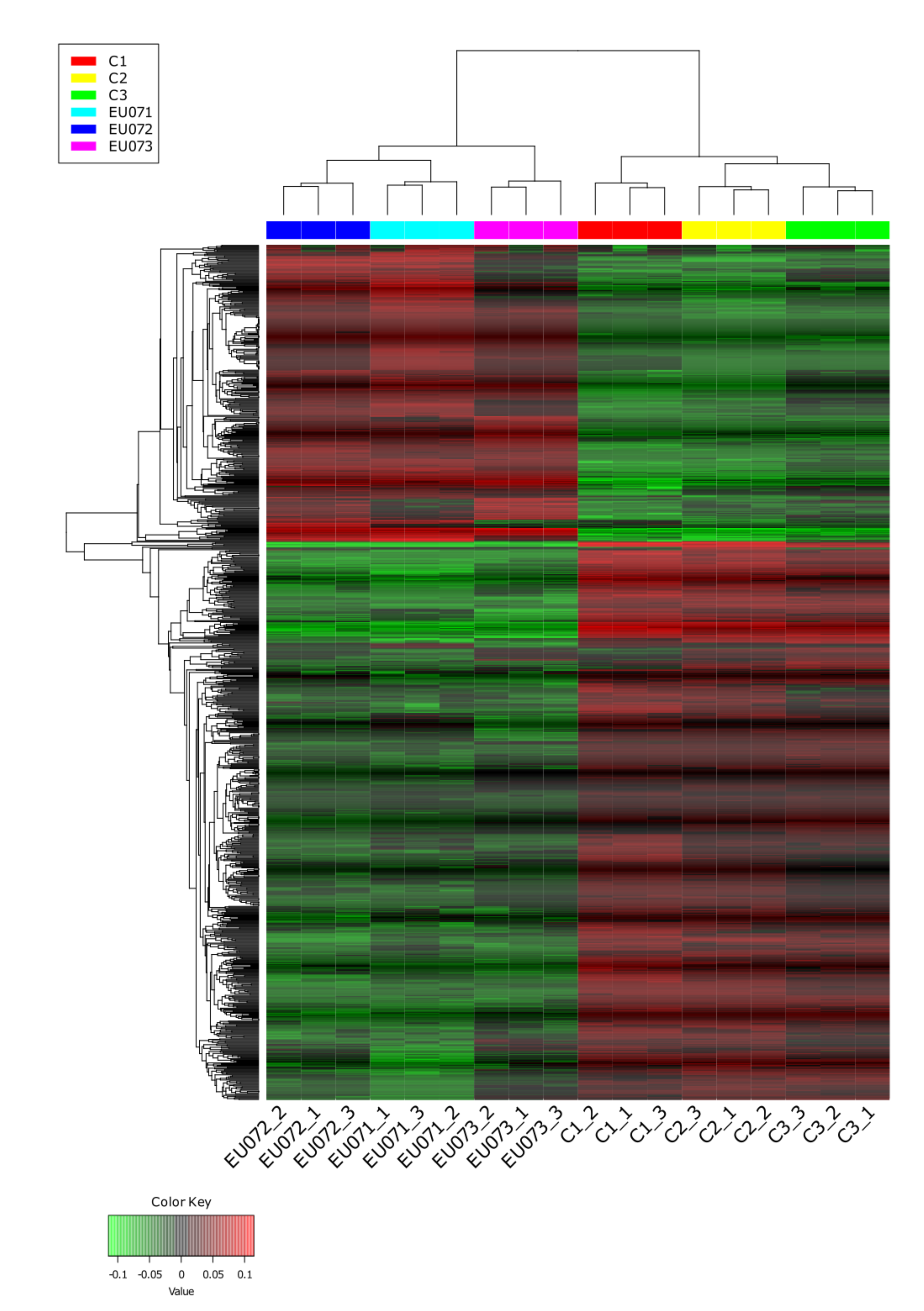
Heatmap of DEGs in *F. graminearum* treated with *B. velezensis* EU07. The heatmap shows genes significantly up-or downregulated in *F. graminearum* following exposure to metabolites secreted by *B. velezensis* EU07. Rows represent genes and columns represent biological replicates. Colour intensity reflects normalized expression levels (red: upregulated, blue: downregulated). Hierarchical clustering highlights co-expression patterns. Only genes with ≥2-fold change and false discovery rate (FDR) < 0.05 are shown.

The 50 most strongly up- and downregulated genes are shown in Figure 3A. Overall gene expression patterns across treatment replicates are summarised as bar graphs (Figure 3B). Categorisation of DEGs by biological process, cellular component, and molecular function revealed logarithmic changes in expression (Table 2). In addition, relationships among DEGs in four process categories were visualised using principal coordinate analysis, presented as heatmaps (Figure 3C).

**Figure 3.**
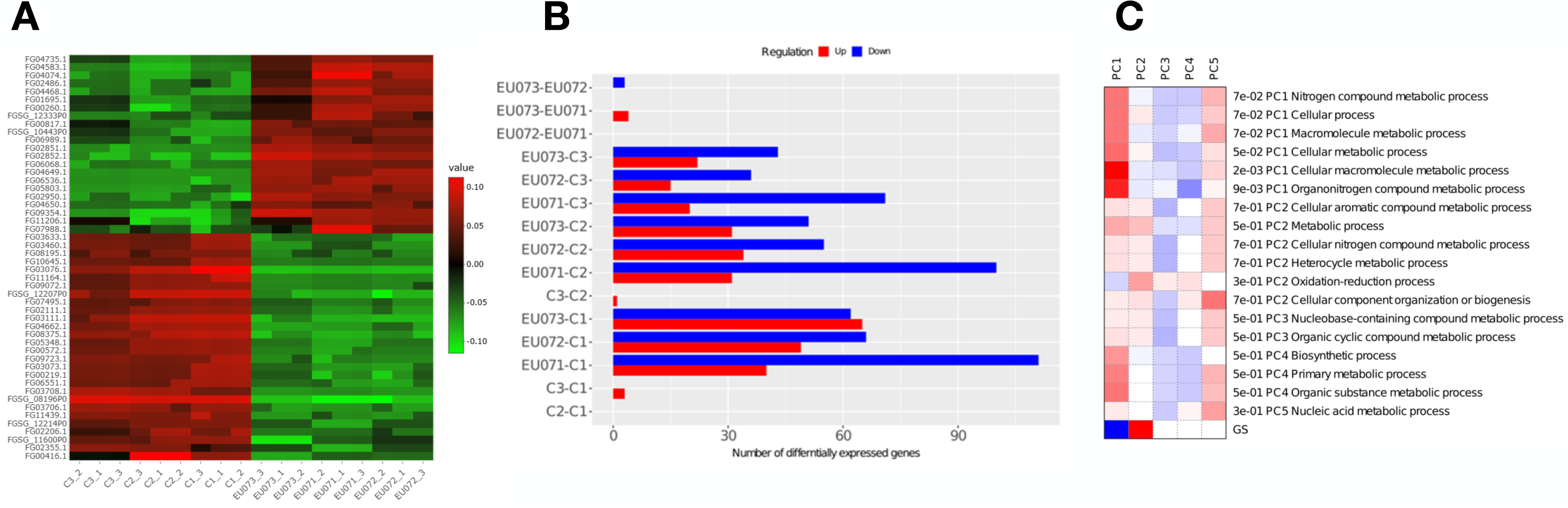
Transcriptomic responses of *F. graminearum* to metabolites of *B. velezensis* EU07. **A)** Heatmap showing the most variably expressed genes in *F. graminearum* following exposure to EU0-derived metabolites. **B)** Bar chart summarizing the number of DEGs, categorized by upregulation and downregulation. **C)** Principal Coordinate Analysis (PCoA) displaying clustering of metabolic processes associated with DEGs in *F. graminearum* upon exposure to EU07-derived metabolites.

**Table 2.**
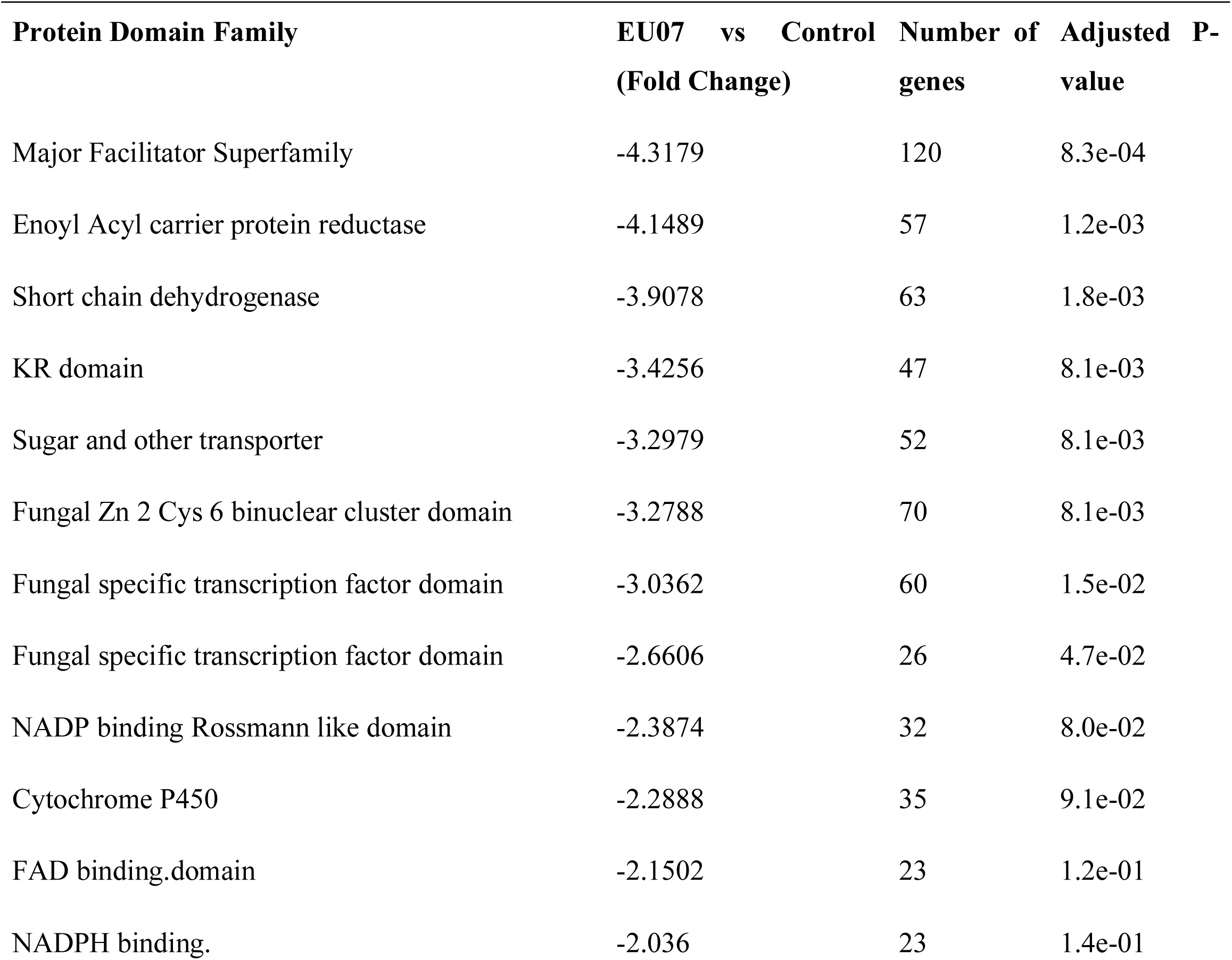
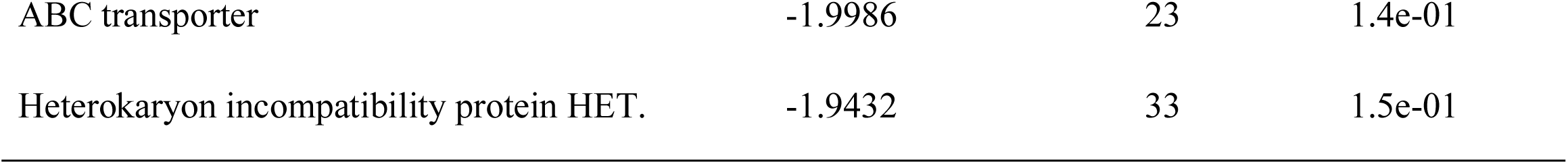
Genes downregulated in *F. graminearum* upon exposure to *B. velezensis* EU07 products and their associated protein families.

| Protein Domain Family | EU07 vs Control<br>(Fold Change) | Number of genes | Adjusted P-value |
| --- | --- | --- | --- |
| Major Facilitator Superfamily | -4.3179 | 120 | 8.3e-04 |
| Enoyl Acyl carrier protein reductase | -4.1489 | 57 | 1.2e-03 |
| Short chain dehydrogenase | -3.9078 | 63 | 1.8e-03 |
| KR domain | -3.4256 | 47 | 8.1e-03 |
| Sugar and other transporter | -3.2979 | 52 | 8.1e-03 |
| Fungal Zn 2 Cys 6 binuclear cluster domain | -3.2788 | 70 | 8.1e-03 |
| Fungal specific transcription factor domain | -3.0362 | 60 | 1.5e-02 |
| Fungal specific transcription factor domain | -2.6606 | 26 | 4.7e-02 |
| NADP binding Rossmann like domain | -2.3874 | 32 | 8.0e-02 |
| Cytochrome P450 | -2.2888 | 35 | 9.1e-02 |
| FAD binding, domain | -2.1502 | 23 | 1.2e-01 |
| NADPH binding. | -2.036 | 23 | 1.4e-01 |
| ABC transporter | -1.9986 | 23 | 1.4e-01 |
| Heterokaryon incompatibility protein HET. | -1.9432 | 33 | 1.5e-01 |

### Functional categorization of the top DEGs and enrichment analysis

Gene expression profiling of *F. graminearum* exposed to *B. velezensis* EU07 revealed widespread transcriptional reprogramming. Application of EU07 cells suppressed 111 genes and induced 40 genes among 3355 detected transcripts (Figure 3B). The top most downregulated genes are listed in Table 2. A complete list of DEGs classified by Gene Ontology (GO) category is provided in Supplementary Table S2.

The transcriptional response of *F. graminearum* to EU07 treatment extended beyond passive fluctuations, instead reflecting a coordinated adjustment of functional pathways. Several of the most strongly induced genes encode proteins implicated in extracellular remodeling, transcriptional regulation, transport, secondary metabolism, and detoxification. For instance, FGSG_04649, associated with extracellular component restructuring, was upregulated 6.8-fold, suggesting reinforcement of fungal surface defenses or altered interactions with the external environment. Similarly, FGSG_09354, encoding a zinc finger transcription factor, upregulated 6.6-fold, consistent with a role in regulatory reprogramming of downstream gene networks.

Metabolic remodeling was also evident. FGSG_02852 (glycoside hydrolase) was induced 6.5-fold, potentially contributing to carbohydrate degradation or nutrient acquisition. FGSG_04583, encoding a polyketide synthase, showed a 5.9-fold increase, pointing to activation of secondary metabolite biosynthesis. Transporter genes FGSG_02851 (ABC transporter) and FGSG_04074 (MFS transporter) were induced 5.5-and 5.3-fold, respectively, consistent with altered metabolite trafficking across membranes. In addition, genes involved in detoxification and redox balance, such as FGSG_04468 (cytochrome P450 monooxygenase) and FGSG_06068 (aldo/keto reductase), were upregulated by 5.3- and 5.2-fold. Other genes of less well-characterized function, including FGSG_06536 (l-pipecolate oxidase) and FGSG_05803, also showed notable induction (5.1- and 4.9-fold).

In contrast, several genes were strongly repressed, pointing to targeted downregulation of transport, metabolic, and signalling pathways. FGSG_08196 and FGSG_12519, likely involved in cellular transport and cell wall processes, were reduced by ∼10- and 9.7-fold, respectively. FGSG_03111, annotated as an amino acid permease/mitochondrial carrier protein, was downregulated 8.5-fold. Similarly, FGSG_04662 (oxidoreductase), FGSG_12207 (mitochondrial transporter), and FGSG_08375 (serine protease) showed reductions of 7.9-, 7.6-, and 7.2-fold, respectively, while FGSG_11270 (putative kinase) and FGSG_09072 (calcium-binding protein) were repressed by ∼7-fold. Downregulation extended to transcriptional regulators (FGSG_11164, −7-fold) and membrane-associated proteins (FGSG_11439, −6.9-fold). Two additional genes of unknown function, FGSG_13802 and FGSG_11146, were reduced 6.8- and 6.5-fold.

We further examined five genes (FGSG_08196, FGSG_12519, FGSG_04649, FGSG_09354, and FGSG_02852) with the lowest expression levels and found that they are conserved across multiple *Fusarium* species (Supplementary File 1).

Together, these patterns indicate that EU07 products elicit a dual strategy in *F. graminearum*: strong induction of genes associated with defense, metabolic adaptation, and environmental interaction, accompanied by repression of genes linked to transport, signalling, and energy metabolism. This coordinated reconfiguration highlights an adaptive transcriptional program, balancing the costs of stress responses with the need to maintain cellular homeostasis.

### Gene enrichment and network analysis

Pathway enrichment analysis of DEGs revealed associations with virulence-related functions in *F. graminearum*. Interestingly, downregulated genes were enriched in the RmlC-like cupin domain superfamily, while upregulated genes were enriched in amino acid transporters and permeases (Table 3). These results suggest that EU07 metabolites modulate fungal transcriptional programs by suppressing cupin-domain proteins and enhancing transport-related functions.

**Table 3.**
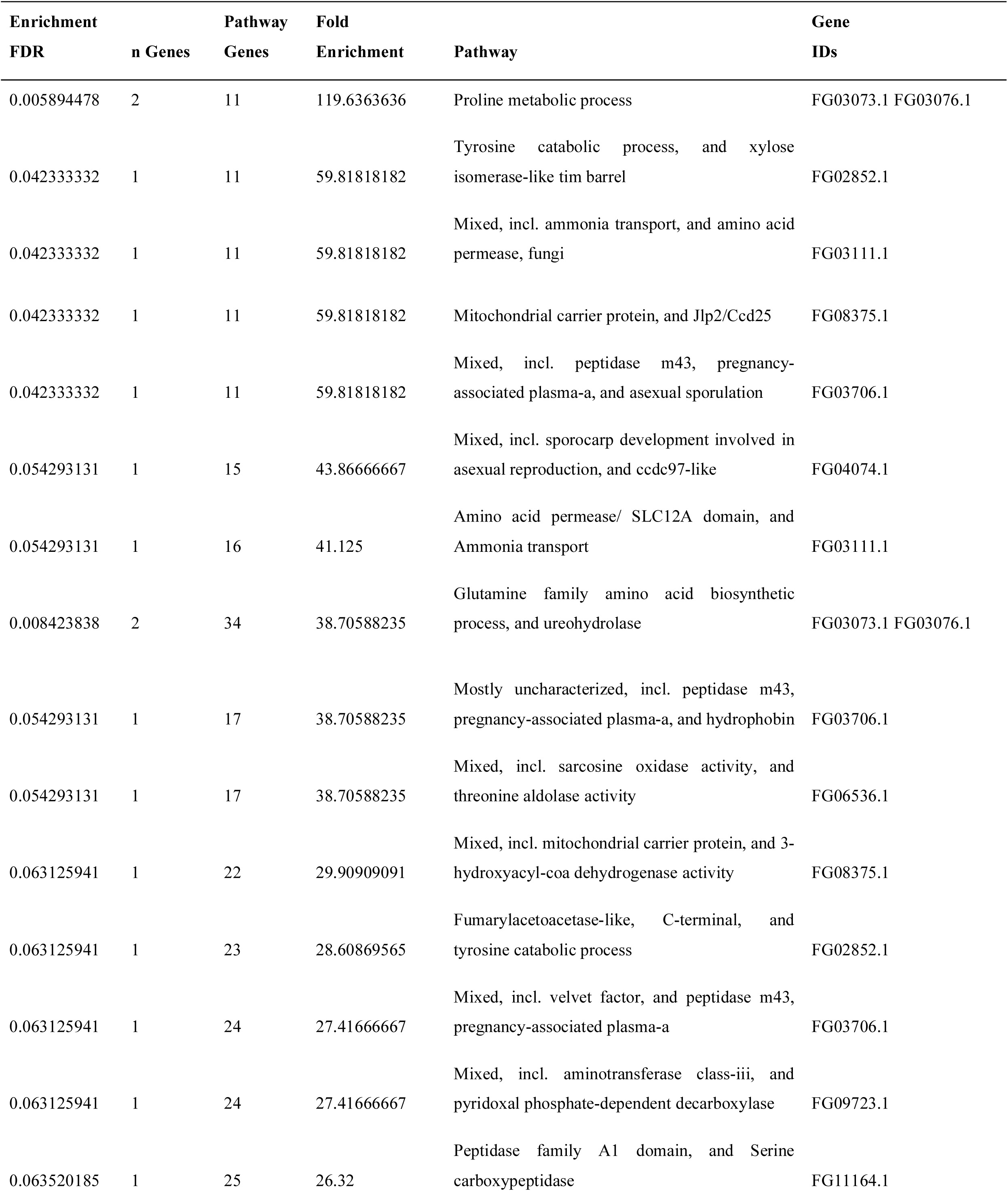
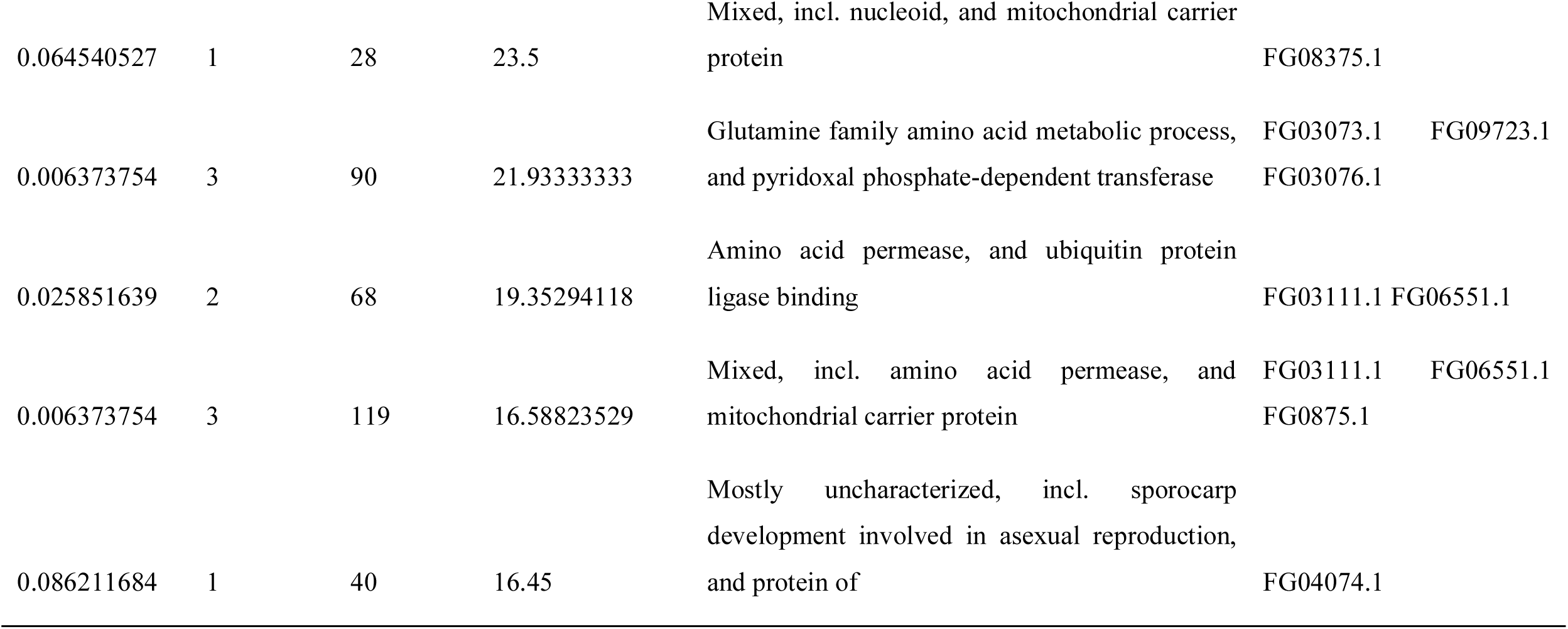
Gene enrichment analysis of *F. graminearum* in response to *B. velezensis* EU07 metabolites and associated functional pathways based on RNA-seq data..

Network analysis using STRING and Cytoscape identified connections between DEGs and functional clusters. The STRING database linked DEGs to KOG1339 and KOG1721, which appear to act cooperatively during the fungal response (Figure 4B). GO classification further supported functional clustering of DEGs within biological process categories (Figure 4A–B; Supplementary File S2).

**Figure 4.**
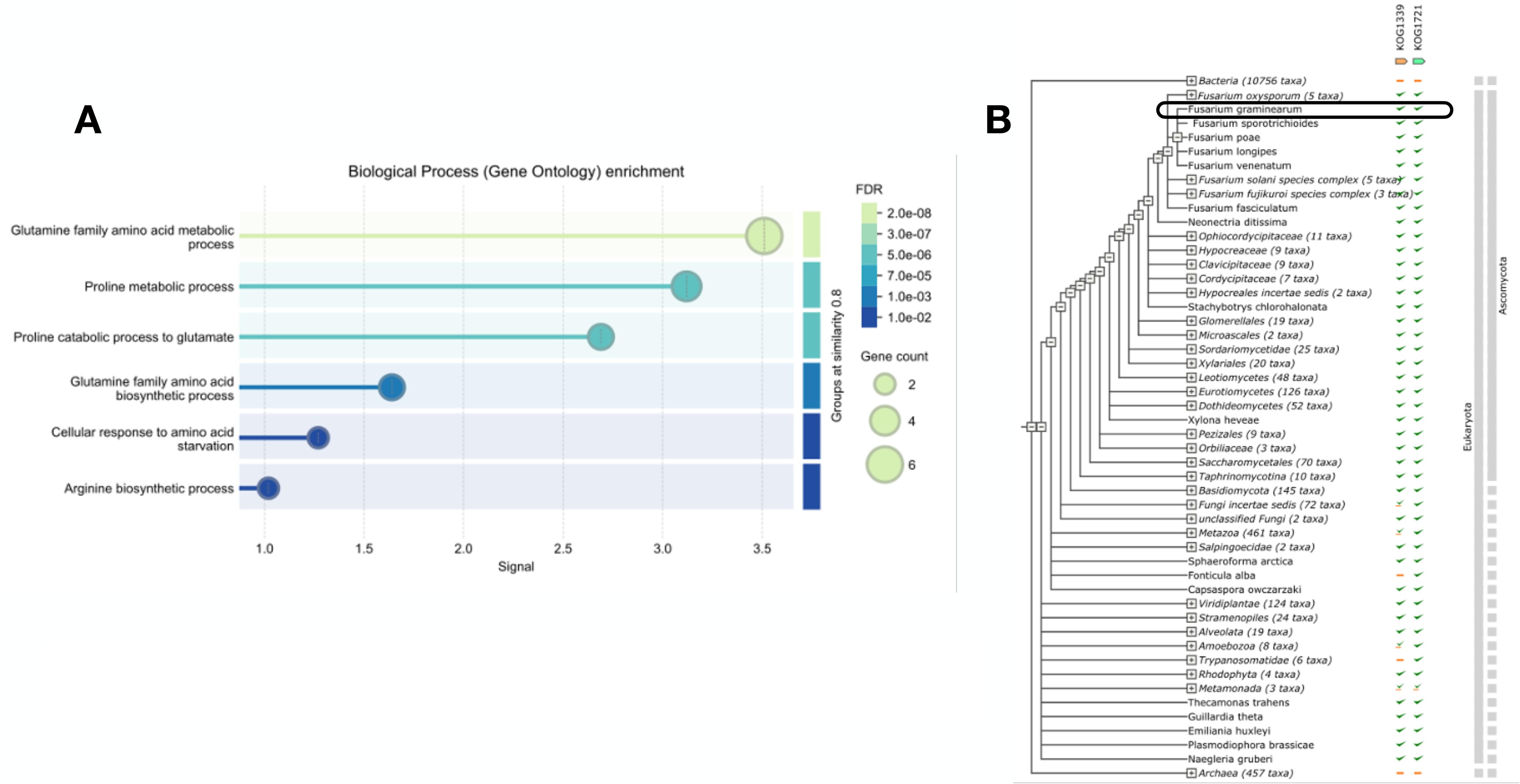
Functional categorization of DEGs in *F. graminearum* in response to *B. velezensis* EU07 metabolites. **A)** GO classification of DEGs associated with biological process. The bar plot displays enriched GO terms, categorized by process type (e.g., metabolic process, cellular process, response to stimulus). Bars are color-coded: blue for upregulated genes, red for downregulated genes. The enrichment significance threshold was set at *p* ≤ 0.05 with a false discovery rate (FDR) < 0.05. The number of genes contributing to each GO category is indicated on top of each bar. **B)** KOG functional classification of DEGs, derived from STRING database annotation and validated by RNA-seq expression data. Functional categories are represented by bars, with color codes indicating expression direction: green for upregulated, orange for downregulated. Statistical significance was assessed using adjusted *p*-values (Benjamini-Hochberg correction, *q* < 0.05).

Integration of RNA-seq data with the STRING database enabled visualization of gene–gene relationships within a network framework related to genes showing down regulation in *F. graminearum* (Figure 5A). Cytoscape analysis confirmed coordinated expression patterns, identifying subsets of genes that interact directly or indirectly. Genes without connections were considered functionally unlinked under the tested conditions (Figure 6B–C). Collectively, these analyses demonstrate that EU07 treatment induces a coordinated yet selective reprogramming of *F. graminearum* transcriptional networks.

**Figure 5.**
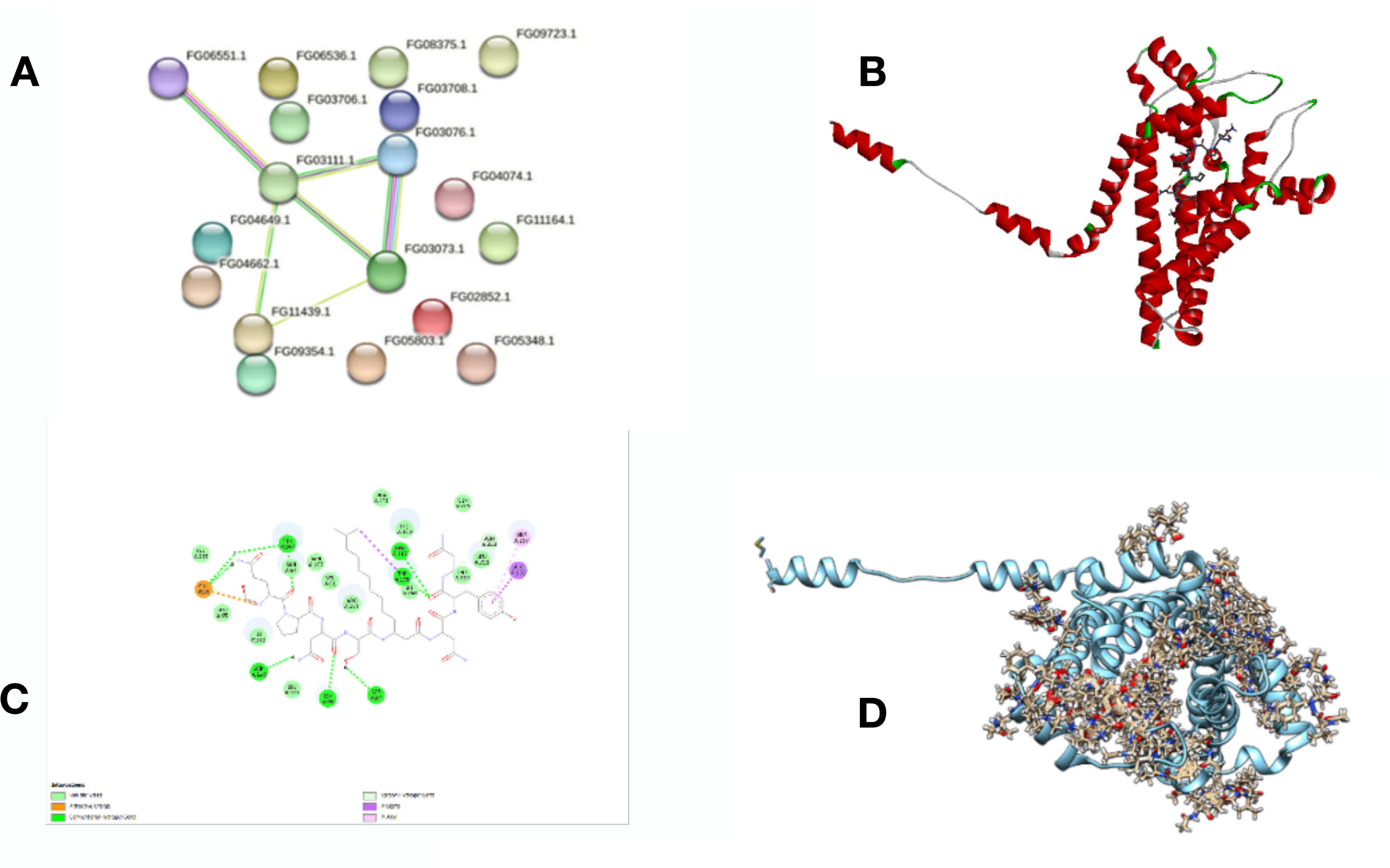
Molecular interaction between *B. velezensis* EU07-derived Iturin and *F. graminearum* virulence-associated proteins. **A)** Gene co-expression network of the top 10 most significantly downregulated genes in *Fusarium graminearum* exposed to EU07 metabolites, indicating hub genes potentially involved in virulence suppression. **B)** 3D molecular docking model showing the binding of Iturin (ligand) to the Apolipophorin protein (receptor). **C)** 2D interaction diagram illustrating the key amino acid residues (GLU68, GLN264, ASP158, LYS60, ASP57, LYS260, TYR108, ARG112, THR204, LEU211, ASN212, ALA208, ARG207) involved in the Iturin-Apolipophorin binding interface. **D)** 3D visulasation of all possible ligand conformations, showing the flexibility and binding hotspot regions of Iturin on the Apoliphorin surface.

**Figure 6.**
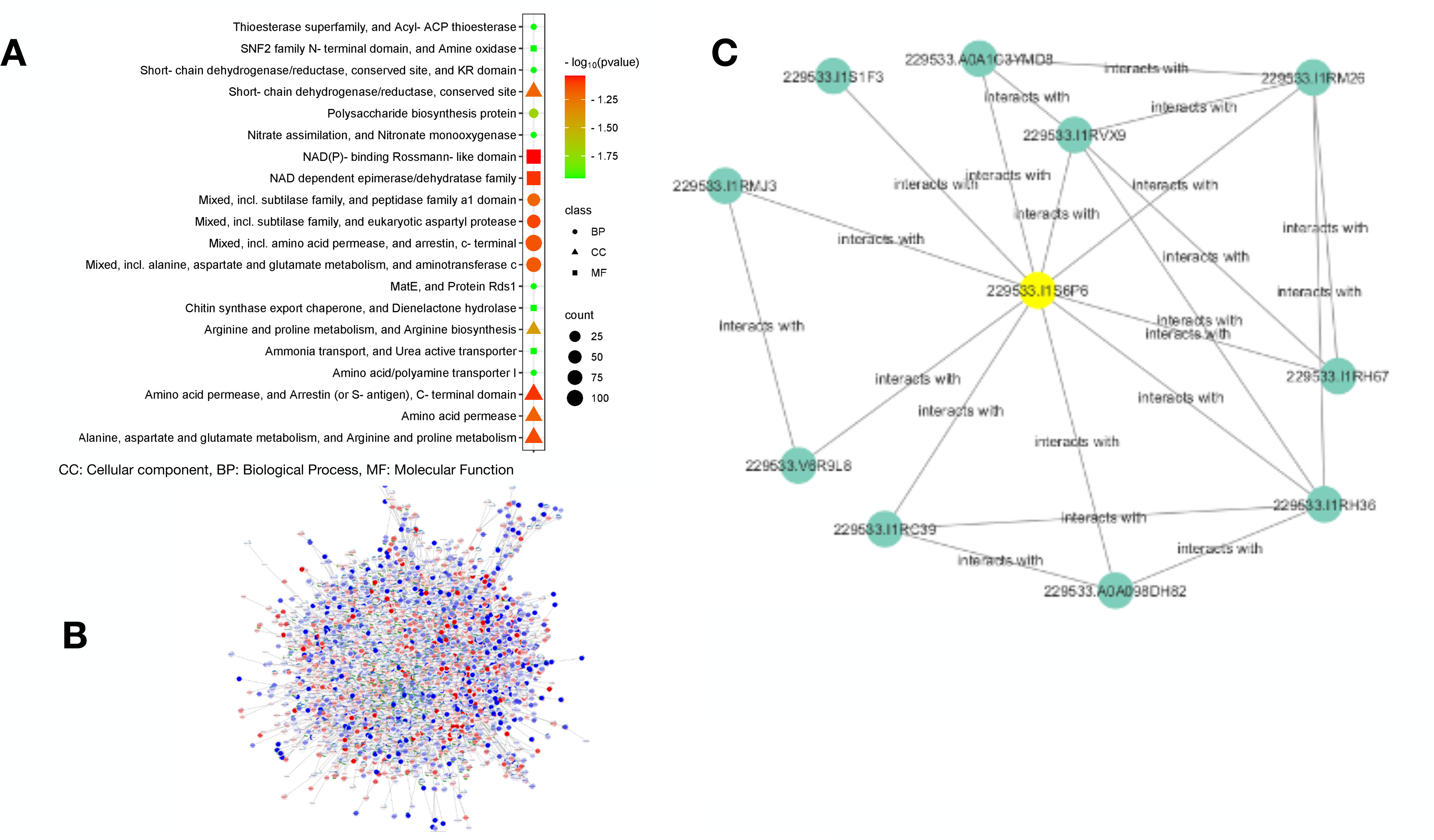
Functional classification and gene network analyses of *F. graminearum* DEGs following exposure to *B. velezensis* EU07 metabolites. **A)** GO classification of DEGs in *F. ggraminearum* following treatment with the *B. velezensis* EU07 metabolites. Categories include Cellular Component (CC), Biological Process (BP), and Molecular Function (MF). Bars represent enrichment significance as – log (P value). **B)** Co-expression network of the most significantly down-regulated genes, illustrating gene clustering and potential functional interactions. **C)** Integrated gene interaction network constructed using RNAseq data (SIF format) and String database annotations, visualized via Cytoscape. The colour changes on each node represents the specific expressed gene folding (up/down regulated) of *F. graminearum* upon exposure to EU07 metabolites.

### Molecular docking studies

RNA-seq analysis identified several hypothetical protein-coding genes in *F. graminearum* with altered expression in response to EU07. Among these, the apolipophorin protein, implicated in virulence and membrane function, was significantly downregulated. Given that EU07 produces the lipopeptide iturin, we performed molecular docking to evaluate potential interactions between iturin and apolipophorin.

Docking simulations predicted a stable interaction with a binding energy of −7.2 kcal/mol (Figure 5B, Supplementary File S3). The complete docking conformation of iturin with apolipophorin, characterized by helical structures, was investigated, focusing multiple iturin compound binding events on the target protein. The highest binding energy between apolipophorin, (receptor) and iturin (ligand) and its coordinates on receptor shown in Figure 5B, Supplementary File S4). The iturin ligand was predicted to interact with multiple residues of apolipophorin, including GLU68, GLN264, ASP158, LYS60, ASP57, LYS260, TYR108, ARG112, THR204, LEU211, ASN212, ALA208, and ARG207 (Figure 5C). All predicted binding poses of ligand on receptor are also shown in Figure 5D. These results suggest that EU07-derived iturin may directly target apolipophorin, potentially compromising fungal membrane-associated virulence functions.

### Gene Ontology (GO) classification and integrated network analysis

To assess the functional consequences of transcriptional changes in *F. graminearum* exposed to *B. velezensis* EU07 metabolites, Gene Ontology (GO) classification and network analyses were performed. DEGs were assigned to Biological Process (BP), Molecular Function (MF), and Cellular Component (CC) categories. Enrichment was visualized based on –log(P value), with significant terms including membrane organization, cellular transport, and oxidoreductase activity (Figure 6A). These categorised point to disruption of fungal homeostasis by bacterial metabolites.

Two complementary network analyses were conducted. First, a co-expression network was generated from the most strongly downregulated genes, revealing tightly clustered nodes associated with virulence, membrane integrity, and stress responses (Figure 6B). This focused view highlights functional connections among repressed genes.

Second, an integrated interaction network was built from the full RNA-seq dataset in SIF format (Supplementary File 2), enriched with STRING interactions and visualized in Cytoscape (Figure 6B). Nodes were coloured according to fold-change direction and magnitude, enabling visualization of both induced and repressed genes (Figure 6B). Filtering highlighted subsets of functionally related DEGs, while unconnected nodes represented genes with no detectable interactions under the tested conditions (Figure 6C).

Together, these analyses provide both a targeted and a global perspective on the transcriptional reprogramming of *F. graminearum* in response to EU07.

## Discussion

Our study demonstrates that exposure of *F. graminearum* K1-4 (*Fg*-K1-4) to *B. velezensis* EU07 induces pronounced morphological and physiological changes. After 48 h of treatment, fungal cultures exhibited swollen hyphae and conglobated structures, in contrast to untreated controls. Comparable stress responses have been reported in filamentous fungi exposed to *Bacillus* strains or their metabolites (Baysal et al., 2013; Gong et al., 2015; Patel et al., 2024).

Lipopeptides such as fengycins, surfactins, and iturins are known to disrupt fungal membranes. Deleu et al. (2008) reported concentration-dependent membrane disruption by these compounds in a DPPC unilamellar vesicle model, with surfactins exhibiting up to 40-fold greater effects than fengycins. Patel et al. (2024) demonstrated that simple forms of fengycins, including agrastatin1 and plipastatin A1, induce pore formation in fungal membranes. Baysal et al. (2013) showed that volatile organic compounds (VOCs) from *Bacillus* strains, including EU07, caused distorted, swollen, and disrupted mycelia of *F. oxysporum*, consistent with our observations. VOC analysis of EU07 identified a protein (ID 3835), which shares 99% identity with SAM-dependent methyltransferases, which are typically involves in the methylation of small molecules critical for metabolism and secondary metabolite biosynthesis (Sun et al., 2020).

Gong et al. (2015) further demonstrated that purified iturin A and plipastatin A treatments caused deformation, lateral expansion, and ultrastructural damage to *F. graminearum* conidia. Our experiments with *Fg*-K1-4 treated with EU07, using both broth and bacterial pellet, produced similar morphological and ultrastructural changes. These findings guided our choice of EU07 pellet treatment for transcriptomic analysis, ensuring that observed gene expression changes reflected direct bacterial–fungal interactions rather than medium-related effects. Sampling at 6 h post-treatment allowed the capture of early transcriptional responses (Schrey et al., 2005; Gu et al., 2017; He et al., 2017).

RNA-seq analysis revealed strong downregulation of genes encoding apolipophorin and proline dehydrogenase (PRODH). Apolipophorin III proteins are exchangeable apolipoproteins that play critical roles in lipid transport, membrane structure, and fungal virulence (Wang et al., 2002). PRODH (EC 1.5.5.2) catalyzes the oxidation of L-proline to Δ¹-pyrroline-5-carboxylate and regulates intracellular proline, an osmoprotectant that contributes to ROS detoxification, mitochondrial protection, and stress tolerance (Rizzi et al., 2017; Ali et al., 2018). Δ¹-pyrroline-5-carboxylate dehydrogenase, a downstream mitochondrial enzyme, also participates in stress mitigation and immune defense. The coordinated downregulation of these genes indicates that EU07 metabolites compromise fungal membrane integrity, stress resilience, and metabolic homeostasis.

Molecular docking supported this mechanism, indicating that EU07-derived iturin binds apolipophorin with high affinity (−7.2 kcal/mol), interacting with residues GLU68, GLN264, ASP158, LYS60, ASP57, LYS260, TYR108, ARG112, THR204, LEU211, ASN212, ALA208, and ARG207 (Figure 5B–D). These interactions suggest that EU07 metabolites may directly target membrane-associated virulence factors, consistent with the observed morphological and transcriptomic responses.

Beyond individual genes, RNA-seq profiling showed coordinated upregulation of genes involved in amino acid transport, secondary metabolism, and detoxification pathways, reflecting a multilayered adaptive response to bacterial stress. Functional enrichment and network analyses confirmed that DEGs cluster in pathways related to virulence, membrane organization, transport, and stress responses (Figures 5–6). Collectively, these global patterns highlight the fungal strategy to selectively induce protective pathways while repressing energetically costly functions, such as mitochondrial transport, serine proteases, and kinases, under bacterial challenge.

Spray-induced gene silencing (SIGS, Bilir et al, 2022) has emerged as a promising strategy for managing *F. graminearum* and reducing Fusarium head blight (FHB) severity. Field trials demonstrated that naked aqueous dsRNA sprays targeting core fungal regulatory genes such as *CHS3b* and *MGV1* significantly reduced both FHB incidence and deoxynivalenol (DON) accumulation, with two applications achieving over 90% control in some cases (Feng et al, 2025). Similar success has been reported in controlled environments: dsRNA targeting *TRI6*, a key transcriptional regulator of trichothecene biosynthesis, reduced gene expression, disease spread, and DON accumulation on wheat heads under greenhouse and growth chamber conditions (Hao et al., 2021). Together, these studies confirm the feasibility of SIGS for both disease and toxin suppression.

Our findings extend this field by identifying novel candidate targets that have not yet been explored in SIGS applications. Several strongly downregulated genes, including apolipophorin and *PRODH*, were functionally linked to virulence and stress responses. These genes, revealed through integrated morphological, transcriptomic, and docking analyses, represent previously untested but potentially valuable targets for RNA-based silencing strategies. Importantly, the overlap between genes suppressed by EU07 metabolites and those considered viable SIGS targets suggests complementary avenues for intervention. In line with previous proposals for engineering plants to express dsRNAs or hairpin RNAs (Morozov et al., 2019), our results provide a rational framework for prioritizing functionally critical genes. Coupling such candidate targets with advances in SIGS delivery platforms may accelerate the development of durable, environmentally sustainable RNA-based strategies for FHB and DON management. Potential limitations include the use of broth cultures, which may not fully mimic plant-pathogen interactions, and the early time point (6 h) for transcriptomic sampling, which may not capture longer-term responses. Additionally, some downregulated genes remain hypothetical, requiring functional validation. Future studies should therefore validate candidate targets via gene knockouts or RNAi, assess EU07 effects under plant-pathogen conditions, and evaluate the combined effects of VOCs, lipopeptides, and dsRNA strategies to maximize biocontrol efficacy.

In summary, *B velezensis* EU07 induces profound morphology changes and transcriptional reprogramming in *F. graminearum* K1-4. The upregulation of genes linked to secondary metabolism, transport, and stress adaptation, alongside downregulation of membrane-and metabolism-related genes, highlights a coordinated fungal defense program under bacterial metabolites. Molecular docking results suggest that EU07 lipopeptides may directly target fungal virulence proteins. These insights reveal key mechanisms of bacterial–fungal interaction and identify candidate genes for targeted RNAi-based control, supporting the potential of EU07 as a sustainable biocontrol agent against FHB.

## Material and Methods

### Fusarium graminearum and Bacillus strains

The *F. graminearum* isolate K1-4 (*Fg*-K1-4) used in this study was maintained as previously described by Jimenez-Quiros et al. (2022). The fungus was cultured on potato dextrose agar (PDA) at 22-24°C and was periodically subcultured on spezieller-nährstoffarmer agar (SNA) or on 25% strength PDA to reactivate macroconidia production. The *B. velezensis* strain EU07 used in this study was previously characterized at the genome level by Baysal et al. (2024).

### Dual culture assay for transcriptomics

To determine the optimal conditions for assessing the effect of *Bacillus* EU07 on *F. graminearum*, 50 mL of potato dextrose (PD) broth was inoculated with 100 µL of macroconidia adjusted to 10^6^ conidia/ml and incubated at 24°C with agitation (150 rpm) for two days. Once the fungal cultures exhibited homogeneous growth, they were treated with either sterile H₂O (control) or *Bacillus* EU07. Briefly, bacterial broths were grown for 24 hours at 28°C (OD_600_ of 1), and 4 mL aliquots were centrifuged at 4000 rpm for 10 minutes to obtain a pellet. The supernatant was discarded, and the pellet was resuspended in 4 mL of sterile H₂O before being added to the fungal cultures. Sterile H_2_O was used as control. The bacterial-fungal interaction was allowed to proceed for 6 hours. Following incubation, 2 mL samples were collected from each flask, transferred into cryogenic tubes, flash-frozen in liquid nitrogen, and stored at -80°C for further analysis. Three independent biological replicates per treatment were used (six flasks in total) (Van den Berge et al., 2019).

### RNA isolation and sequencing

Fungal mycelium treated with control and *Bacillus* was ground in liquid nitrogen using a mortar and pestle. Total RNA was isolated using TRIzol reagent (Invitrogen, UK) according to the manufacturer’s instructions. RNA integrity was assessed using an Agilent 2100 Bioanalyser. RNA samples with RIN ≥7 from treated and untreated *Fg*-K1-4 were sequenced at Novogene (UK) on an Illumina platform, with poly-A-captured cDNA libraries (250–300 bp inserts) and paired-end reads (Q30 ≥ 80%). Quality control was conducted at all stages including sample assessment, library preparation, and sequencing. The resulting raw reads were used for subsequent analyses.

### Differential Gene Expression Analysis

The paired-end reads for each sample were imported into the Galaxy software platform (The Galaxy Community, 2022) and mapped to the reference genome of *F. graminarium* (NC_026474.1) using default parameters, except for adjustment of the maximum insert size of the paired-end library. FastQ data were first processed with optimized trimmomatic tools (adjusted parameters for MINLEN, LEADING, CROP and HEADCROP settings) to remove adaptor sequences and low-quality bases (Chen 2023). Subsequently, RNA STAR was used to generate BAM files from pair-end sequences after MultiQC analysis and then visualized using Tablet (Hutton Institute ver. 1.21.02.08; Milne et al., 2011). Raw count data from multiple samples were retrieved and merged into a single matrix using *featureCounts*. Differential expression analysis was performed using the *limma-voom* pipeline in R (via Sambomics tools) to estimate gene expression changes and log2 fold-change (logFC) values. The data were submitted to the iDEP platform for visualization of heatmaps and other gene count-based analyses (Ge et al., 2020). To identify WP values corresponding to gene IDs, BLAST analyses were performed. For gene annotation, BLASTx searches (E-value < 1e-3) were conducted against the NCBI nr database using unigenes as queries. The BLAST results were then imported into Blast2GO for GO term assignment and functional categorization (Conesa et al., 2005). Genes were considered differentially expressed if they exhibited a fold change of ≥2 or ≤−2, with a false discovery rate (FDR)-adjusted *p*-value ≤ 0.05. The protein sequences of five genes with the lowest expression levels were retrieved from the GenBank database. BLASTp analyses were then performed using the *Fusarium* taxid to determine whether these genes are conserved across other *Fusarium* species. A phylogenetic tree was constructed using the neighbour-joining method. Gene enrichment analysis was performed using ShinyGO v0.741 (Ge et al., 2020) with the *F. graminearum* STRINGdb as the reference, and results were visualized accordingly.

### Enrichment gene interaction map and network analysis

An enrichment map was generated from RNA-seq output data using ShinyGO (Ge et al., 2020), an open-source platform for functional enrichment and network visualization. In the enrichment map pathways were represented as nodes, with edges representing shared genes between pathways, thereby illustrating functional relationships. To investigate gene-gene interactions both within and across pathways, selected genes were annotated using the STRING database (http://string-db.org; Szklarczyk et al., 2011). Interaction networks were constructed and visualised in Cytoscape (Shannon et al., 2003). RNA-seq expression values and corresponding .SIF files were integrated into Cytoscape to visualize and validate interaction networks.

### Molecular docking-based virtual screening for protein-ligand interactions

The experimental X-ray diffraction structure of Iturin A and its corresponding protein model, identified as Apolipophorin based on FASTA sequence translation into amino acids using ExPASy (https://www.expasy.org/; Berman et al., 2003), were used for docking studies. Missing residues were added with PyMOL’s builder plugin (v2.5.0), and loop regions containing these residues were refined using MODELLER (v10.1) (Webb and Sali, 2016; PyMOL Molecular Graphics System, 2023). The structure was further processed by removing all heteroatoms except those associated with cofactors, introducing polar hydrogen atoms as required, and assigning Kollman charges (Morris et al., 2009).

A grid box (17 Å × 24 Å × 24 Å) was defined to cover the predictive active site of the modelled structure. Virtual screening was then performed to assess interactions between the Iturin A ligand and the target protein within the predefined grid. Docking simulations were conducted with an exhaustiveness level of 64 using AutoDock Vina (v1.1.2) (Trott and Olson, 2010). Ligands with the highest binding affinity scores were selected for further evaluation under the same docking configuration.

### Protein-ligand interaction profiling

The best docking pose of the ligand was loaded with the simulated protein structure into PyMOL, and all residues within 4 Å of the compound were visualized to identify potential hydrophobic interactions, hydrogen bonds, and ionic interactions. Predicted interactions were cross-validated using TU Dresden’s Protein–Ligand Interaction Profiler (PLIP) webserver, and only interactions confirmed by both manual inspection and PLIP analysis were considered (Salentin et al., 2015).

## Accession numbers

The datasets generated in the current study are available under the BioProject accession number PRJNA1322080.

## Author contributions

MT conceived the study and, with CJQ, designed the experiments. CJQ conducted laboratory work, ÖB performed bioinformatic analyses, and BCK revised the manuscript. All authors contributed to writing and approved the final version.

## Conflict of Interest

The authors declare that there is no conflict of interests.

## Funding

CJQ was funded by the University of Worcester, and the authors gratefully acknowledge the BBSRC partnering award BB/X018253/1 awarded to MT.

## Data availability statement

The data that support the findings of this study are available from the corresponding author on reasonable request. All genomic data are publicly available as described in the paper.

## Supplemental Materials

**Supplementary Table S1. Summary of RNA-seq sequencing reads and quality metrics.**

**Supplementary Table S2: DEGs obtained from the RNA-seq analysis.**

**Supplementary File S1. Phylogenetic trees of five genes showing conservation across *Fusarium* species.**

**Supplementary File S2. Protein–Protein Interaction Network Data in SIF Format**

**Supplementary File S3. Molecular docking model generation details.**

**Supplementary File S4. Structural and Sequence Information for Uncharacterized Protein A0A2H3GNL7 from *Fusarium graminearum***

## Notes

### Competing Interest Statement

The authors have declared no competing interest.

