## Supplementary File S1 for "*Bacillus velezensis* EU07 suppresses *Fusarium graminearum* via transcriptomic reprogramming": Supplementary File S1.pdf

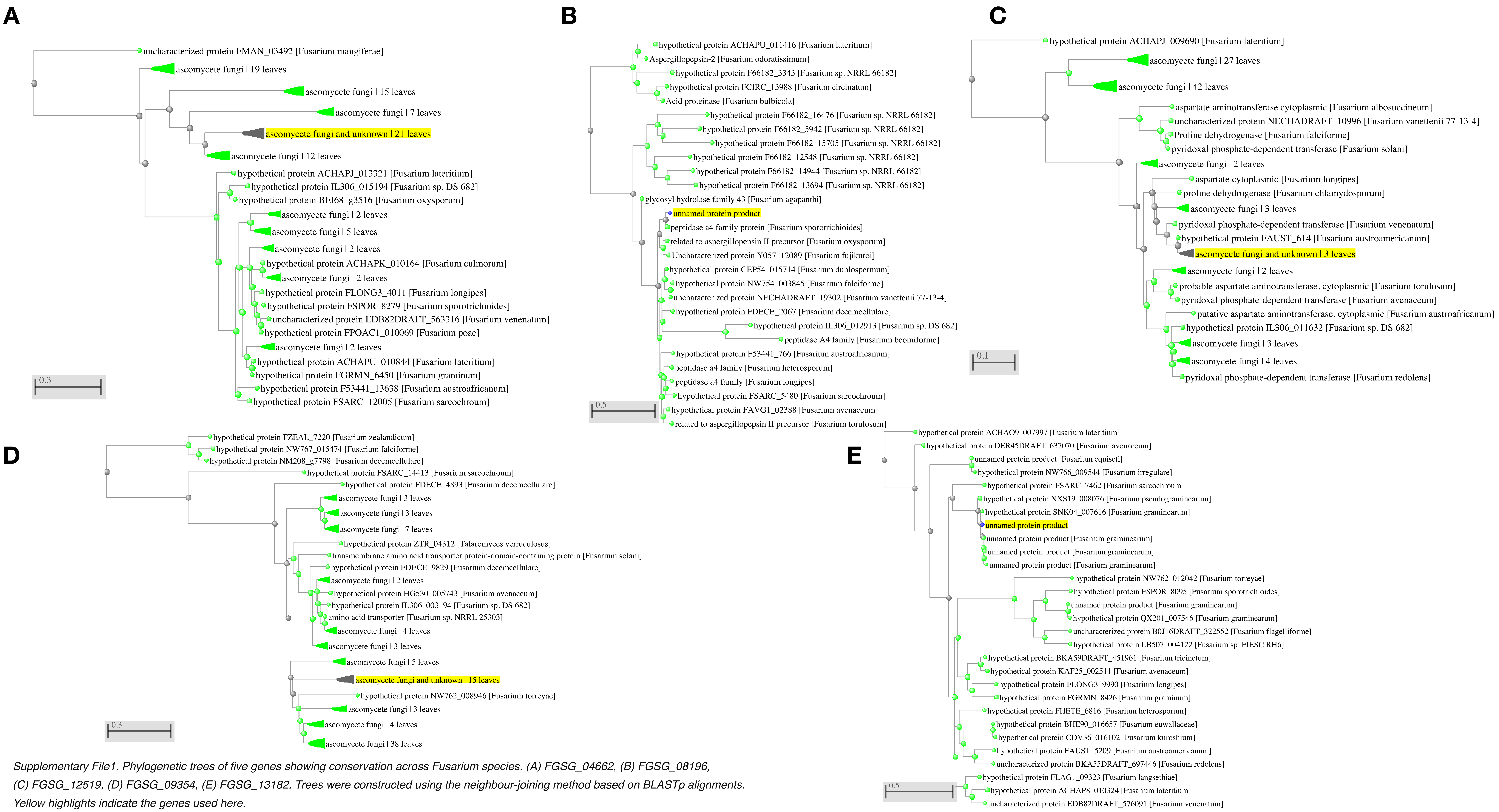


*Supplementary File1. Phylogenetic trees of five genes showing conservation across Fusarium species. (A) FGSG_04662, (B) FGSG_08196, (C) FGSG_12519, (D) FGSG_09354, (E) FGSG_13182. Trees were constructed using the neighbour-joining method based on BLASTp alignments. Yellow highlights indicate the genes used here.*
