## Supplementary File S4 for "*Bacillus velezensis* EU07 suppresses *Fusarium graminearum* via transcriptomic reprogramming": Supplementary File S4.pdf

Search SWISS-MODEL Repository

### A0A2H3GNL7 (A0A2H3GNL7_GIBZA) *Gibberella zeae (Wheat head blight fungus) (Fusarium graminearum)*

#### Uncharacterized protein

347 aa; Sequence (Fasta)

50 100 150 200 250 300

A0A2H3GNL7

monomer 1-347

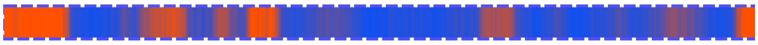

No models have been built for this target sequence.

Build Models

#### Available Structures

#### 1 AlphaFold Model

| | Model ID | Oligo-state | Avg pLDDT | Range | Trg-Mdl Seq Id (%) |
|---|---|---|---|---|---|
| | AF-A0A2H3GNL7-F1 | monomer | 76.50 | | 100.0 |
| | *Download* | | Compare | | *Structure Assessment* |

#### Alignments

#### AF-A0A2H3GNL7-F1-model_v4

```
     Target MRSHAGPITWASVVWLVLGLFQLANCASVDNKLSHGRERI  40
AlphaFold.A MRSHAGPITWASVVWLVLGLFQLANCASVDNKLSHGRERI  40

     Target WLWEMYNIFVDIEGADKQDLILPQDKNAKFVYRRHYSLDK  80
AlphaFold.A WLWEMYNIFVDIEGADKQDLILPQDKNAKFVYRRHYSLDK  80

     Target NTVDGRLSYAEFMADLEAKLTPLDMDYSLQAPNSEGIVPG 120
AlphaFold.A NTVDGRLSYAEFMADLEAKLTPLDMDYSLQAPNSEGIVPG 120

     Target KDGQASGKPSVAEAVETLKLKGWAQTMKVDQVTSGKTKDY 160
AlphaFold.A KDGQASGKPSVAEAVETLKLKGWAQTMKVDQVTSGKTKDY 160

     Target GELVGRVDKKFQDYYDKLGSDPKWKRKLIETNQRTNKLTG 200
AlphaFold.A GELVGRVDKKFQDYYDKLGSDPKWKRKLIETNQRTNKLTG 200

     Target EIVALRQQEADDWVIKQLTQDADHNNPKVPDEDKKLGIGL 240
AlphaFold.A EIVALRQQEADDWVIKQLTQDADHNNPKVPDEDKKLGIGL 240

     Target SRDDLVIERVRSSIPGATTYERVDLVRTFLEHPNQAEFLE 280
AlphaFold.A SRDDLVIERVRSSIPGATTYERVDLVRTFLEHPNQAEFLE 280

     Target KLSTAGIKGGIQGLINWADMLGNERYANWGAGFTDANKSH 320
AlphaFold.A KLSTAGIKGGIQGLINWADMLGNERYANWGAGFTDANKSH 320

     Target HQAKSLWLAMHDSSTKRLGGLSPSCSR              347
AlphaFold.A HQAKSLWLAMHDSSTKRLGGLSPSCSR              347
```

**AlphaFold** AF-A0A2H3GNL7-F1-model-v4; Created: 2022-06-01;
Average Model Confidence (pLDDT) : 76.50 ; Local Confidence

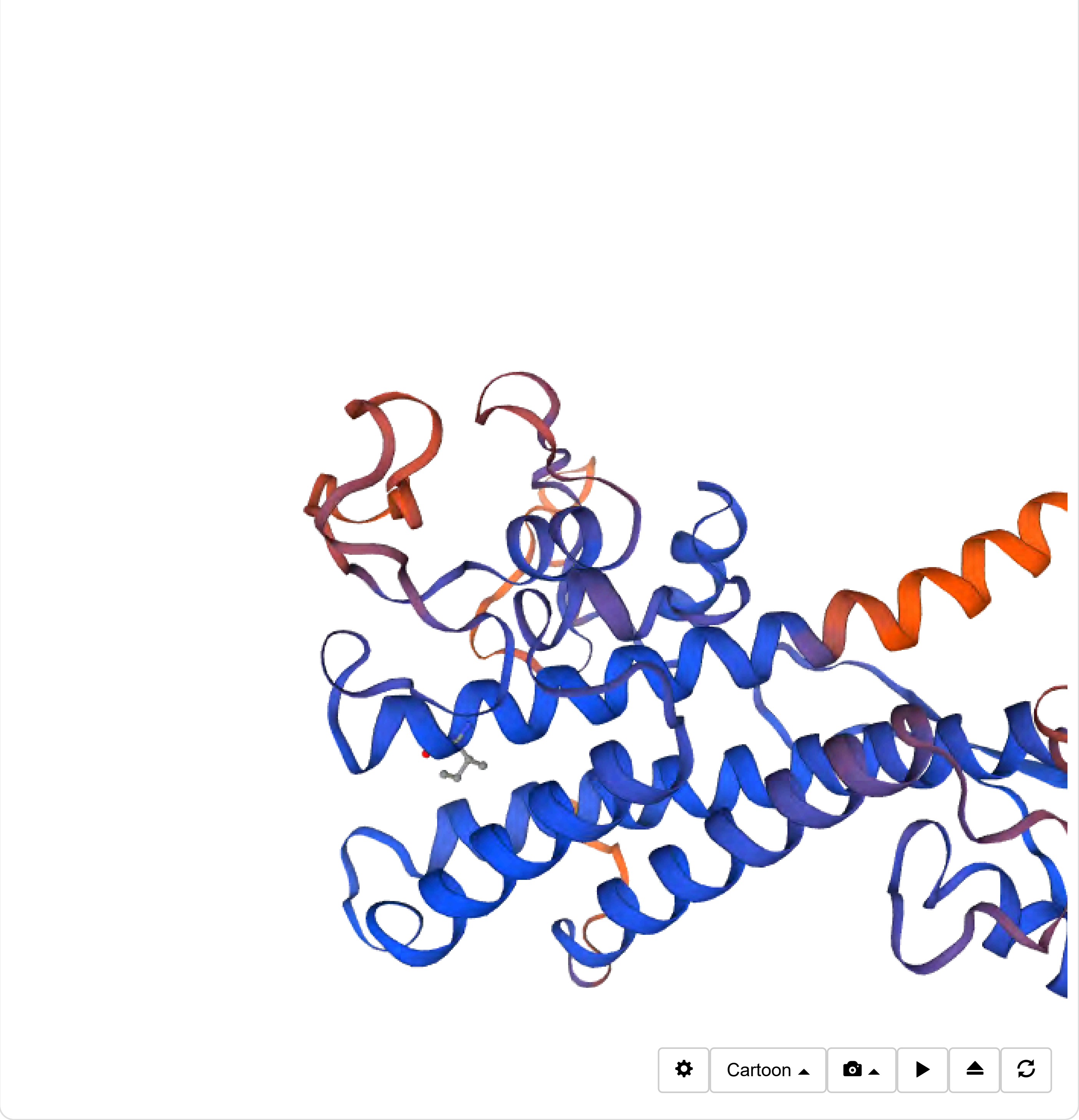

Cartoon
